# NF1 splicing reprograms ERα signaling to promote luminal breast cancer progression

**DOI:** 10.64898/2026.05.21.726647

**Authors:** Patrick S. Dischinger, Ian Beddows, Sophia Agrusa, Marie Adams, Emily Wolfrum, Carrie R. Graveel, Matthew R. Steensma

## Abstract

Alternative splicing is an emerging driver of cancer progression, yet its role in modulating tumor suppressor function remains incompletely understood. Here, we identify an alternatively spliced *NF1* isoform lacking the nuclear localization sequence (NLS; NLS SE) as a clinically and functionally relevant regulator of breast cancer progression. Analysis of TCGA and AURORA cohorts revealed that *NF1* NLS SE is enriched in metastatic tumors, preferentially occurs in luminal subtypes, and is associated with decreased overall survival independent of *NF1* genomic alterations.

Using a CRISPR-engineered MCF7 model, we show that NLS SE expression abolishes nuclear localization of neurofibromin, enhances ERK signaling, and promotes proliferation and resistance to endocrine and MAPK-targeted therapies. Despite increased ERα protein levels, transcriptomic analysis revealed suppression of canonical estrogen response programs and activation of KRAS, EMT, and inflammatory pathways. Mechanistically, NLS SE expression increased RNA-bound ERα and reprogrammed RNA binding protein networks, including CELF, ESRP1, and SRSF family members, leading to widespread alternative splicing.

Together, these findings define NF1 NLS exon skipping as a luminal breast cancer-associated, isoform-level mechanism that disrupts nuclear neurofibromin function, rewires ERα signaling towards post-transcriptional regulation, and promotes endocrine-resistant disease. Targeting splicing regulatory networks may represent a therapeutic strategy in *NF1-*dysregulated breast cancer.

Graphical Abstract.
NF1 NLS exon skipping drives cytoplasmic retention of neurofibromin and reprograms ERα toward post-transcriptional regulatory functions.
(A) In cells expressing NLS-containing NF1 isoforms, neurofibromin localizes to the nucleus and constrains ERα activity. (B-C) Alternative splicing of the NF1 NLS exon generates the NLS SE isoform, resulting in cytoplasmic retention of neurofibromin. This shift disrupts canonical ERα signaling and is associated with increased ERα phosphorylation and enhanced RNA-binding activity. ERα engages RNA and cooperates with splicing machinery, including phosphorylated SF3B1, to promote transcriptomic remodeling. These changes support a therapy-resistant state characteristic of aggressive, endocrine-resistant breast cancer.

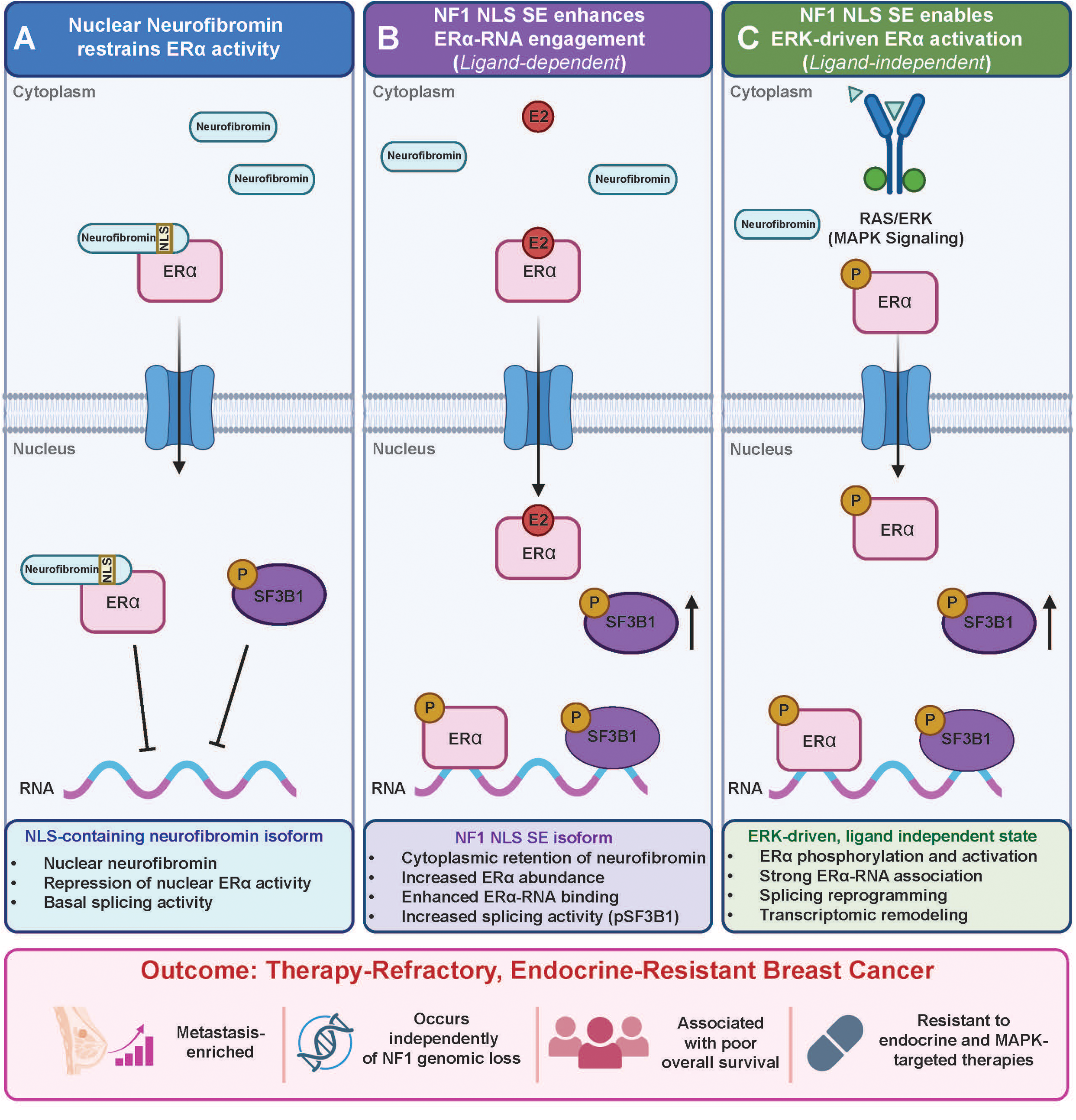

## Introduction

Breast cancer remains the most commonly diagnosed malignancy in women worldwide and remains a major health challenge due to the complexity of its pathogenesis and diverse clinical presentations, which complicate prevention and have profound effects on identifying appropriate treatment efforts. With incidence rates continuing to rise, particularly in the United States where approximately 321,910 new cases of invasive breast cancer are projected to be diagnosed in women in 2026, a comprehensive understanding of breast cancer molecular mechanisms and genetic risk factors is essential in developing precision therapies to effectively target varying molecular drivers in disease progression^1^. One of the molecular mechanisms that is relatively understudied in breast cancer is alternative splicing (AS). Despite growing recognition, the contribution of AS to cancer biology remains incompletely understood relative to our understanding of the transcriptomics and genomics, a critical process where RNA-binding proteins (RBPs) generate multiple mRNA isoforms from a single gene^2,3,5^. AS plays a pivotal role in expanding the proteomic diversity of cells, occurring in 92-96% of human coding genes^6,7^. When dysregulated, splicing events in tumor suppressor and oncogene transcripts can result in aberrant protein expression or function, promoting oncogenesis, disease progression, and therapy resistance^4,8,9^. For instance, dysregulated splicing events occur as a result of mutated or dysregulated expression of cancer-related genes such as TP53 and MYC and influence tumorigenic pathways, particularly in pancreatic and breast cancers^10,11^. In breast cancer, up to 50% of tumors exhibit amplifications or overexpression of RBPs, which drive metastasis through the AS of key genes that regulate proliferation and invasion^12,13^. Recently, estrogen receptor alpha (ERα) was identified as a non-canonical RBP, highlighting the intricate and understudied role of AS in cancer biology^14,15^. These insights underscore the need to further investigate the contribution of transcript variants in key oncogenic pathways, such as RAS and ER signaling, to breast cancer progression. Importantly, alternative splicing alterations frequently arise independently of genomic mutations, representing an orthogonal layer of oncogenic regulation that remains underexplored in breast cancer^16,17^.

The *NF1* gene is a *bona fide* driver of breast cancer^18,19^. The *NF1* gene encodes neurofibromin, a large multidomain protein that negatively regulates RAS signaling. Germline *NF1* mutations in the autosomal dominant tumor predisposition syndrome, Neurofibromatosis type I (NF), place women at a substantial risk for breast cancer-related death. Women with NF are at a tenfold increased risk of developing breast cancer under the age of 40^20,21^. Furthermore, NF1 mutations are frequently observed in metastatic breast cancer, suggesting a role for neurofibromin in promoting epithelial-to-mesenchymal transition (EMT)^22^.

Despite the established link between NF1 mutations and breast cancer, we have a limited understanding of the role of NF1 alternative splicing in tumor progression^23^. *NF1* produces at least five distinct and well characterized transcript isoforms through the splicing of exons 9a, 10a-2, 31, 43, and 52^24,25^. Exon 31 encodes an alternative exon within the gap-related domain, where its inclusion or skipping produces the NF1 type II and type I transcript isoforms, respectively. The type I isoform exhibits a stronger binding affinity to RAS, enhancing GTPase activity and promoting negative RAS regulation, however in contrast, the type II isoform has a tenfold lower affinity for RAS, leading to prolonged RAS activity^26,27^. Due to RAS’s oncogenic role in cancer, type I and type II isoforms have been extensively characterized^28–33^.

Emerging evidence suggests that the nuclear localization of neurofibromin, which is strictly governed by exon 52 through a nuclear localization sequence (NLS), also plays a significant role in cancer. The loss of nuclear neurofibromin through alternative splicing leads to the cytoplasmic retention of neurofibromin, preventing it from carrying out its nuclear functions^34^. The nuclear function of neurofibromin has not been fully elucidated. Recently, neurofibromin was demonstrated to function in the nucleus as a co-repressor of ERα transcription both through both physical binding of ERα and other indirect effects^35,36^. The NLS skipped exon isoform (NLS SE) lacks exon 52 and is highly expressed during development in several rat tissues, but its expression declines with age, suggesting its crucial role in development^37^. Here, we show that NLS SE expression is not only associated with breast cancer disease progression, but also altered ERα function and rewired splicing networks to promote cancer cell plasticity and therapeutic resistance.

Given the importance of *NF1* mutations in breast cancer and the emerging recognition of neurofibromin nuclear functions, there is a critical need to define how *NF1* transcript variation contributes to disease progression. In this study, we investigated the impacts of *NF1* NLS SE using integrated analysis of TCGA and AURORA breast cancer cohorts together with a CRISPR-engineered MCF7 model.

We identify NLS SE as a recurrent, metastasis-enriched splicing event that occurs independently of *NF1* genomic loss, is preferentially associated with luminal tumors, and predicts poor clinical outcome. Functionally, NLS SE abolishes nuclear neurofibromin localization, enhances ERK and ERα signaling, and promotes resistance to endocrine and MAPK-targeted therapies, Mechanistically, loss of nuclear neurofibromin shifts ERα from canonical transcription regulator to a non-genomic, RNA-associated factor that cooperates with RNA-binding proteins to drive widespread splicing reprogramming.

These findings establish *NF1* alternative splicing as a distinct regulatory axis in breast cancer and identify NLS SE as a clinically relevant mechanism linking nuclear neurofibromin loss to endocrine resistance and disease progression.

## Results

### Alternative Splicing of NF1 Nuclear Localization Sequence (NLS) is Enriched in Breast Cancer Disease Progression

Given the known role of *NF1* genetic mutations in driving breast cancer progression, we aimed to determine how alternative *NF1* splicing impacts RAS and ER signaling. We interrogated both the TCGA and AURORA breast cancer datasets which includes primary breast tumors and matched metastatic tumors across all PAM50 subtypes. We conducted replicate Multivariate Analysis of Transcript Splicing (rMATS) to compare differential splicing events between primary and metastatic tumors, as well as identify *NF1* splicing alterations that are associated with disease survival and outcome (Figure 1A)^38^.

**Figure 1.**
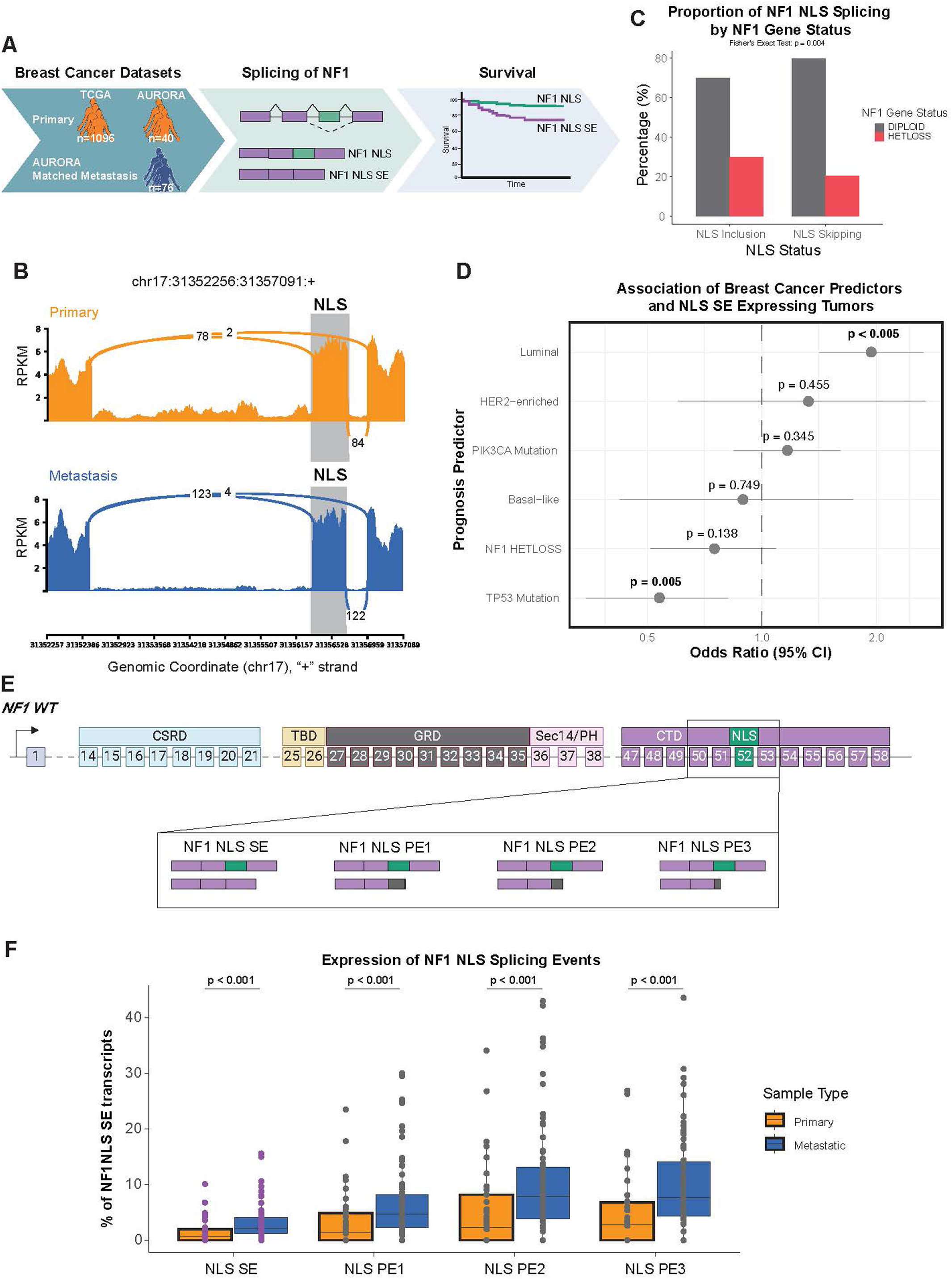
NF1 NLS exon skipping is an independent, luminal-enriched splicing event associated with metastatic progression. (A) Overview of RNA-seq datasets analyzed using rMATS (FDR < 0.05) to identify *NF1* alternative splicing events in primary tumors (TCGA and AURORA) and matched metastatic breast tumors (AURORA). Significant events were subsequently correlated with molecular subtype and patient survival. (B) Sashimi plots illustrating representative exon junction coverage in matched primary and metastatic breast tumors showing increased skipping of the NLS exon (exon 52) in metastatic lesions. (C) Comparison of *NF1* NLS exon skipping (SE) and inclusion across tumors stratified by *NF1* copy number status in the TCGA cohort. Fisher’s exact test revealed enrichment of NLS SE events in diploid tumors relative to those with heterozygous *NF1* loss (HETLOSS) (*p* = 0.004; odds ratio [OR] = 1.65; 95% CI, 1.15–2.39), indicating that NLS exon skipping occurs independently of *NF1* genomic loss. (D) Forest plot of multivariable logistic regression assessing associations between *NF1* NLS SE expression and molecular features, including *NF1* status, *TP53* and *PIK3CA* mutation status, and PAM50 subtype. NLS SE expression was positively associated with luminal subtype (OR = 1.94; *p* < 0.005) and negatively associated with *TP53* mutation (OR = 0.54; *p* = 0.005), while *NF1* copy number loss, HER2-enriched and basal-like subtypes, and *PIK3CA* mutation were not significant (*p* > 0.1 for all). (E) Schematic of *NF1* gene structure highlighting four alternative splicing events within the NLS region identified by rMATS (FDR < 0.05). One event (NLS SE) maintains the open reading frame, whereas three (NLS PE1–3) introduce premature termination codons predicted to trigger nonsense-mediated decay (NMD). (F) Quantification of NLS exon skipping across primary and metastatic samples using a beta-binomial generalized linear mixed model (GLMM) with patient-level random intercepts. All four events showed increased exon skipping in metastases (*p* < 0.001). For the canonical NLS SE event, mean exon skipping was significantly higher in metastatic tumors (2.69%, 95% CI, 1.92–3.75%) than in primary tumors (1.03%, 95% CI, 0.64–1.66%), representing a 2.66-fold increase in odds (*z* = 4.58; *p* < 0.0001).

Our analysis revealed differential splicing events affecting *NF1* transcripts in metastatic tumors compared to their matched primary counterparts. Among the differential splicing events, we observed a significant increase of exon 52 skipping, which encodes the NLS of neurofibromin. Exon 52 is 123 base pairs and its exclusion results in a reading frame shift and loss of NLS, referred to as NLS SE in this study. This skipping event was detected at a 2-fold higher rate in metastatic tumors than in primary tumors (FDR < 0.05), indicating that NLS SE in *NF1* promotes breast cancer disease progression (Figure 1B). Given the established association between *NF1* genomic alterations and poor prognosis in breast cancer, we next examined whether NF1 NLS SE events are linked to underlying *NF1* mutations. Using exon-level splicing data from TCGA SpliceSeq, tumors were classified as exhibiting either NF1 NLS Skipping (NLS SE) or NLS Inclusion (no exon skipping detected)^39^. *NF1* mutation status were obtained from cBioPortal^40^. Fisher’s exact testing revealed that NLS SE events were significantly more frequent in tumors with diploid *NF1* compared to those with heterozygous loss (HETLOSS) (Figure 1C; p=0.004, odds ratio = 1.65, 95% CI: 1.15-2.39). This finding indicates that NF1 NLS SE is not merely a secondary consequence of genomic loss but represents an independent post-transcriptional regulatory mechanism impacting protein localization.

To determine whether NF1 NLS SE is associated with other molecular or clinical breast cancer features, we performed multivariable logistic regression including *NF1* copy number status, *TP53* and *PIK3CA* mutation status, and PAM50 molecular subtype. Consistent with our univariate analysis, *NF1* HETLOSS was not significantly associated with NLS SE (odds ratio [OR] = 0.75, p = 0.138). In contrast, the luminal PAM50 subtype showed a strong independent association with increased odds of NLS SE (OR = 1.94, p < 0.005). Conversely, *TP53* mutation was significantly associated with NLS inclusion (OR = 0.54, p = 0.005), whereas *PIK3CA* mutation and HER2-enriched or basal-like subtypes showed no significant association (all p > 0.3) (Figure 1D). Collectively, these findings indicate that NF1 NLS SE represents a distinct post-transcriptional regulatory event preferentially enriched in luminal breast cancers and independent of canonical *NF1* genomic loss or *TP53/PIK3CA* mutational status. In addition to NLS SE, our rMATS analysis identified three other alternative splicing events involving partial NLS SE, resulting in poison exons (PE1, PE2, and PE3). All four events involved splicing of the NLS of NF1 and revealed a marked increase in exon skipping in metastatic samples, indicating robust depletion of the NLS exon in this context. (FDR < 0.05; > 5 junction reads) (Figure 1E). While the NLS SE splicing event preserved the open reading frame and produced an NLS-deficient, translatable isoform, the remaining three generated premature stop codons (Figure 1E). PEs like ones seen here are known to trigger nonsense-mediated decay and have been implicated in the post-transcriptional silencing of tumor suppressor genes such as *TP53* and *ARID1A*^41,42^. To confirm that these *NF1* splicing events were not driven by outlier patients with multiple metastatic samples, we employed a beta-family generalized linear mixed model (GLMM) with patient-level random effects. This analysis quantified the proportion of NF1 NLS SE transcripts across matched primary-metastatic pairs. All four events demonstrated significantly increased NLS skipping in metastases (p < 0.001) (Figure 1F). Specifically, for the canonical NLS SE event, metastatic tumors exhibited a mean exon-skipping rate of 2.69% (95% CI: 1.92-3.75%) compared to 1.03% (95% CI: 0.64-1.66%) in primaries, corresponding to a 2.66-fold increase in the odds of exon exclusion (z = 4.58, p < 0.0001). Together, these results indicate that metastatic progression is characterized by a preferential enrichment of neurofibromin isoforms lacking the NLS and by the increased generation of non-translatable *NF1* transcripts, suggesting a mechanism for reduced nuclear neurofibromin function in advanced disease.

### NF1 NLS SE is Enriched in ER-Positive Breast Cancers and Associated with Decreased Overall Survival

To assess whether NF1 NLS SE impacts prognosis similar to NF1 mutations, we analyzed the TCGA breast cancer dataset, which includes 1,096 sporadic primary breast tumors. We classified tumors into two groups: those that expressed NF1 NLS SE and those that did not (NLS SE vs. NLS). The PAM50 classification provides a well-established and biologically relevant framework that accounts for breast cancer heterogeneity and contains a robust clinical metadata set. This subtyping method has been extensively validated across diverse cohorts and is widely used in both clinical and translational research to stratify breast tumors based on gene expression patterns^43,44^. Large scale genomic analyses like The Cancer Genome Atlas (TCGA) have demonstrated that PAM50 subtypes are powerful in predicting patient prognosis alongside therapeutic efficacy. Among PAM50 subtypes, 495 primary tumors were available for analysis. In a multivariable Cox proportional hazards model adjusting for mutation burden, age at diagnosis, and NF1, *TP53*, and *PIK3CA* mutational status, patients with tumors expression NF1 NLS SE exhibited significantly reduced overall survival compared to those in the NLS exon inclusion (hazard ratio [HR] = 0.48, 95% CI = 0.28 – 0.82, p = 0.01). Kaplan-Meier analysis confirmed that NF1 NLS SE expression correlated with worse overall survival (OS) (p = 0.01; Figure 2A). Divergence between survival curves emerged at approximately 60 months, a clinically relevant inflection point to the onset of endocrine resistance and metastatic recurrence in ER-positive disease, suggesting that NF1 NLS SE may predict treatment-refractory progression^45,46^.

**Figure 2.**
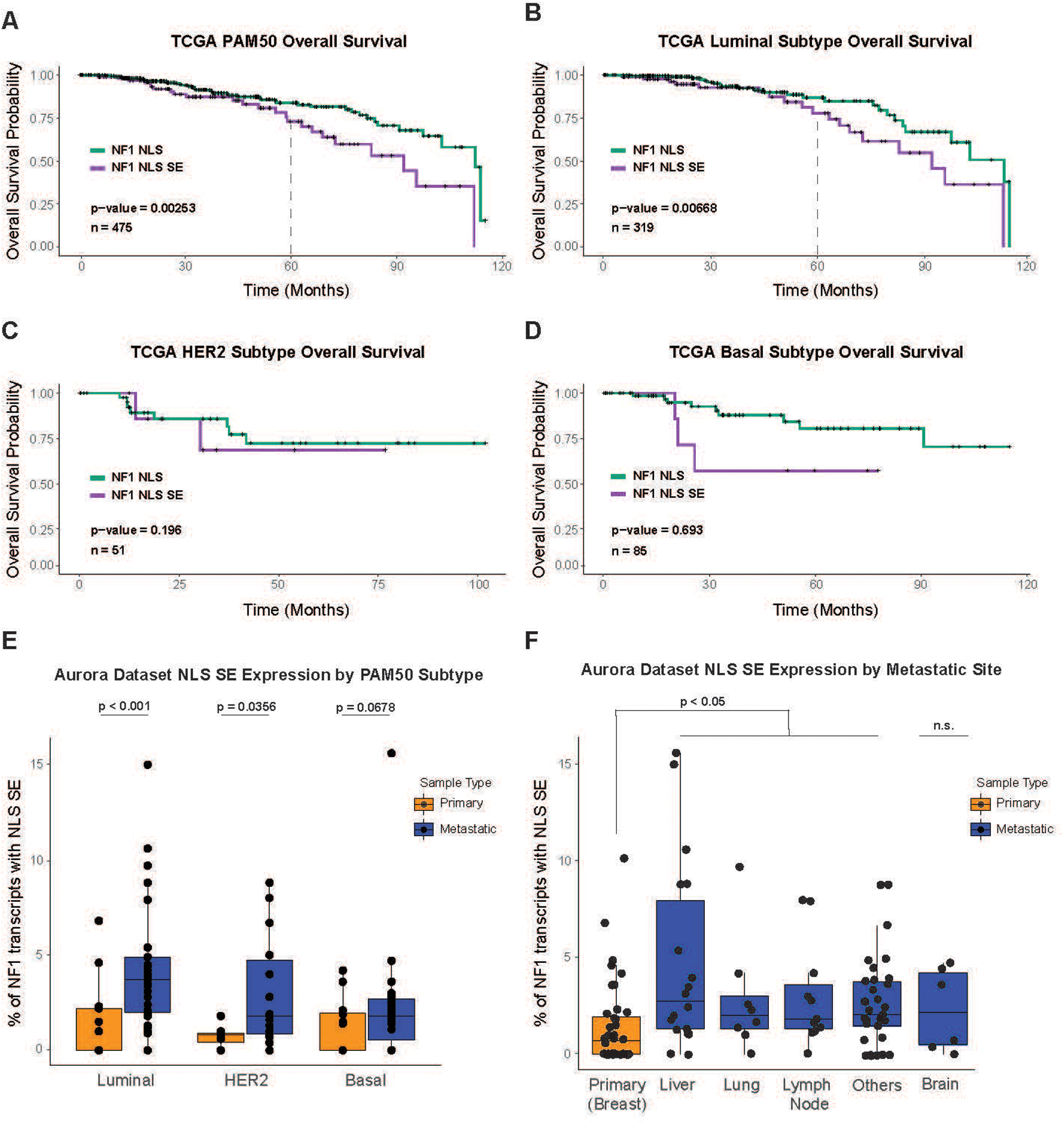
NF1 NLS exon skipping is associated with poor prognosis and metastatic enrichment in luminal breast cancer. (A) Kaplan–Meier overall survival analysis of TCGA breast cancer patients (*n* = 475) stratified by *NF1* NLS splicing status (NLS inclusion vs. NLS skipping, SE). Patients with NLS SE–expressing tumors had significantly reduced survival (log-rank *p* = 0.01), with divergence of survival curves at approximately 60 months, coinciding with the clinical onset of endocrine resistance in ER⁺ disease. Multivariable Cox proportional hazards modeling, adjusted for age at diagnosis, mutation count, and *NF1*, *TP53*, and *PIK3CA* mutation status, confirmed that NLS SE was independently associated with worse survival (HR = 0.48; 95% CI, 0.28–0.82; *p* = 0.00253). Proportional hazards assumptions were validated using Schoenfeld residuals. (B) Subtype-specific Cox analysis in luminal tumors (*n* = 319) showed a significant association between *NF1* NLS SE expression and reduced overall survival (HR = 2.08; 95% CI, 1.22–3.57; *p* = 0.00668), independent of *NF1* genomic loss. Survival curve divergence emerged after ∼60 months, suggesting a link to late-onset endocrine resistance. (C) In HER2-enriched tumors (*n* = 51), a nonsignificant trend toward reduced survival was observed in NLS SE–expressing cases (*p* = 0.196), likely due to limited sample size and event rate. (D) In basal-like tumors (*n* = 85), no significant survival difference was detected between NLS SE and inclusion groups (*p* = 0.693), consistent with the aggressive course and high censoring rate typical of this subtype. (E) Quantification of *NF1* NLS SE expression in primary versus metastatic tumors stratified by PAM50 subtype in the AURORA cohort. A beta-family generalized linear mixed model (GLMM) with patient-level random intercepts was used to compare groups. Luminal metastases showed significantly higher mean NLS SE expression (4.1%; 95% CI, 2.7–6.1%) than primaries (0.68%; 95% CI, 0.32–1.46%), corresponding to a 6.2-fold increase in exon skipping (*z* = 4.92; *p* < 0.001). HER2-enriched tumors exhibited a modest but significant increase (*p* = 0.0356), whereas basal-like tumors showed a nonsignificant trend (*p* = 0.0678). (F) *NF1* NLS SE expression across metastatic sites in the AURORA cohort. Enrichment was observed in liver (*p* = 0.0356), lung (*p* = 0.0678), and lymph node (*p* < 0.05) metastases relative to matched primary tumors, indicating preferential association with sites of distant colonization. NLS SE was quantified as a percentage of total *NF1* transcripts from RNA-seq data.

To further evaluate the prognostic impact of *NF1* splicing within molecular subtypes, we performed subtype-specific Cox regression. In luminal tumors (n = 319), NF1 NLS SE was strongly associated with reduced OS (HR = 2.08, 95% CI = 1.22 – 3.57, p = 0.007) after adjustment for confounding variables (Figure 2B). In contrast, HER2-enriched tumors (n = 51) demonstrated a nonsignificant trend towards poorer survival (p = 0.196; Figure 2C), and basal-like tumors (n = 85) showed no association (p = 0.693; Figure 2D). Although limited by smaller sample sizes, the consistent directional trend across subtypes suggests that *NF1* splicing dysregulation may broadly influence disease progression.

To determine whether NF1 NLS SE represents a metastasis-enriched isoform event, we analyzed RNA-seq data from the AURORA cohort and stratified tumors by PAM50 subtype^47^. Using a beta-family GLMM with patient-level random intercepts, we quantified the proportion of NF1 NLS SE events. In luminal tumors, metastatic samples displayed a significantly higher mean NLS SE level (4.1%, 95% CI = 2.7 – 6.1) compared to primary tumors (0.68%, 95% CI = 0.32 – 1.46), corresponding with a 6.2-fold increase in the odds of exon skipping (z = 4.92, p = < 0.0001; Figure 2E). HER2-enriched tumors exhibited a modest but significant increase (OR = 3.02, p = 0.035), whereas basal-like tumors showed a nonsignificant trend (OR = 1.79, p = 0.067).

To assess whether enrichment was anatomically specific, we next stratified AURORA tumors by metastatic site. GLMM analysis revealed significantly elevated NF1 NLS SE expression in liver, lung, and lymph node metastases compared to matched primary tumors (p < 0.005; Figure 2F). Collectively, these findings demonstrate that NF1 NLS SE is associated with poor prognosis in luminal breast cancer and is preferentially enriched in metastatic lesions, particularly ER-positive tumors. This supports a model in which loss of NF1 NLS through aberrant splicing contributes to endocrine resistance and metastatic adaptation by diminishing nuclear neurofibromin function.

### NF1 NLS SE Enriched Tumors Have Decreased ER-alpha Transcriptional Activity

To investigate how loss of neurofibromin nuclear localization influences metastatic progression, we analyzed primary and metastatic breast tumors from the AURORA cohort and quantified NF1 NLS SE using rMATS. For each patient, we calculated the proportion of *NF1* transcripts containing either NLS inclusion or NLS SE and stratified cases by directionality of splicing change between primary and metastatic lesions. Tumors exhibiting increased NF1 NLS SE expression in metastases were classified as “NLS SE High,” whereas those with decreased NLS SE expression were designated “NLS SE Low” (Figure 3A). This analysis identified 39 NLS SE High and 15 NLS SE Low metastatic tumors, enabling investigation into how altered *NF1* isoform balance shapes transcriptional programs associated with advanced disease.

**Figure 3.**
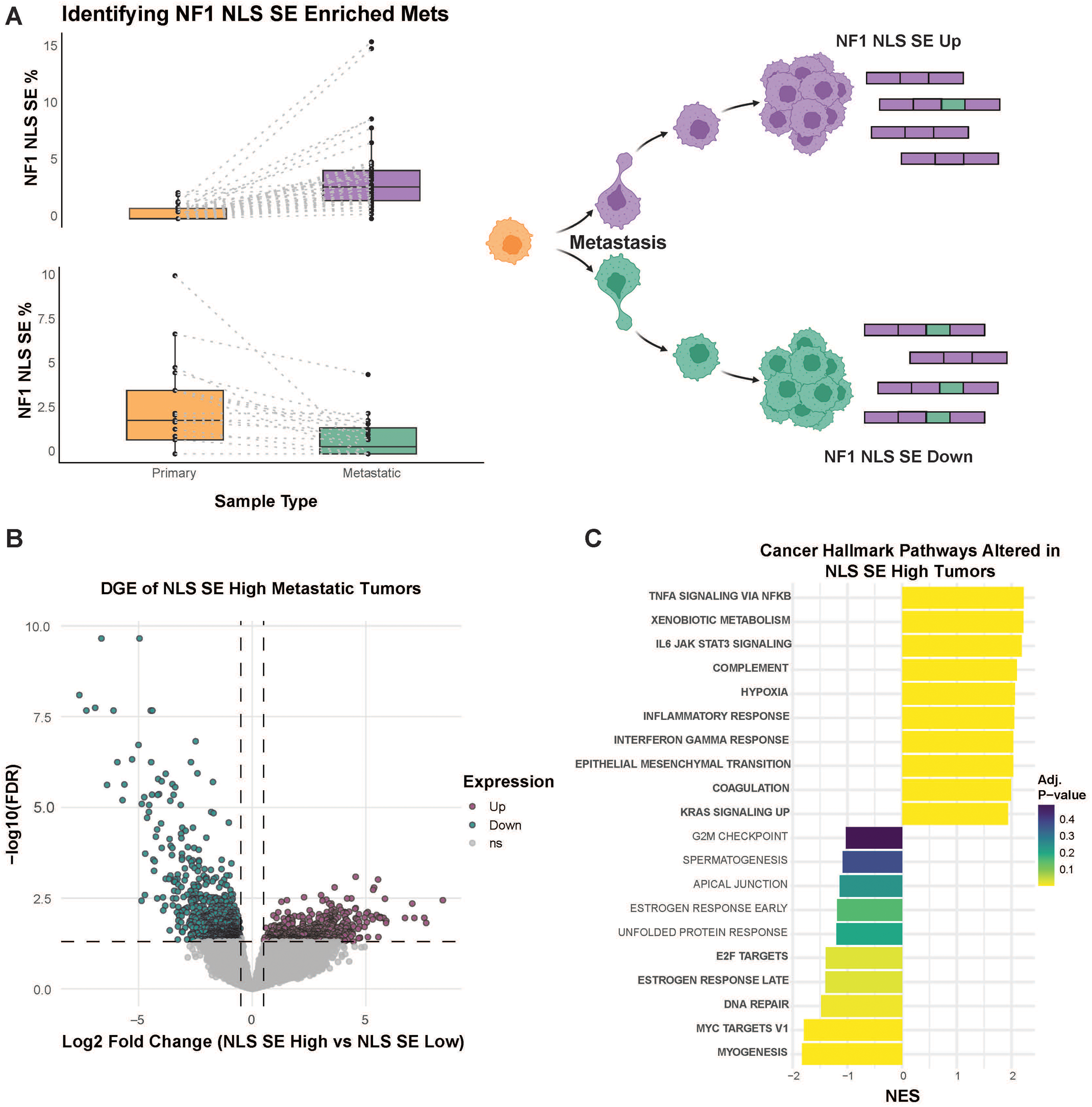
NF1 NLS exon skipping in metastatic tumors induces transcriptomic reprogramming toward mesenchymal and inflammatory phenotypes. (A) Splicing analysis of *NF1* NLS exon skipping was performed using rMATS on matched primary and metastatic breast cancer samples from the AURORA cohort. Patients were stratified based on the direction of splicing change between tumor pairs. Metastatic tumors with increased *NF1* NLS skipping relative to the primary were classified as “NLS SE High” (*n* = 39), whereas those with decreased skipping were classified as “NLS SE Low” (*n* = 15). This grouping enabled transcriptomic comparison according to inferred nuclear neurofibromin abundance. (B) Volcano plot of differentially expressed genes (DEGs) between NLS SE High and NLS SE Low metastatic tumors. A total of 1,243 DEGs were identified (FDR < 0.05; |log₂FC| > 1), including 515 upregulated and 728 downregulated genes in the NLS SE High group. (C) Gene Set Enrichment Analysis (GSEA) of Hallmark pathways revealed that NLS SE High tumors were enriched for epithelial–mesenchymal transition (EMT), KRAS signaling, and TNF-α signaling via NFκB, while *Estrogen Response Late*, *E2F Targets*, and *MYC Targets* pathways were downregulated. These findings indicate that loss of nuclear neurofibromin promotes activation of mesenchymal and pro-inflammatory programs while suppressing ERα-dependent transcriptional activity.

Differential gene expression (DGE) analysis comparing NLS SE High versus NLS SE Low metastases identified 1,243 significantly altered genes (FDR < 0.05, |log_2_FC| ≥ 0.5), including 515 upregulated and 728 downregulated transcripts (Figure 3B). Gene Set Enrichment Analysis (GSEA) of Hallmark pathways revealed that NLS SE High tumors were characterized by upregulation of epithelial-mesenchymal transition (EMT), KRAS signaling, and TNF-alpha signaling via NFκB, consistent with metastatic and inflammatory phenotypes (Figure 3C). In contrast, Estrogen Response Late, E2F Targets, and MYC Targets were significantly downregulated, supporting that loss of nuclear neurofibromin disrupts ER-alpha transcriptional regulation and cell-cycle control.

Collectively, these findings reveal that metastatic tumors enriched for the NF1 NLS SE isoform undergo a transcriptional shift toward mesenchymal and proliferative escape programs while losing estrogen-responsive signaling. This further defines that NF1 NLS SE promotes endocrine resistance and metastatic adaptation.

### *In Vitro* MCF7 Model Demonstrates NF1 NLS SE Expression Drives Enhanced ER-Alpha Signaling and Endocrine Resistance

To investigate further the functional impact of NF1 NLS SE, we generated an in vitro model of NLS loss in the ER-positive breast cancer cell line MCF7 (Figure 4A). Using CRISPR/Cas9 genome editing, we targeted the 5’ end of the NF1 NLS exon to promote exon skipping. The resulting mutant clone, MCF7-C33, expressed both NF1 NLS-included and NF1 NLS SE transcripts, while the empty vector (EV) control expressed only the canonical NLS-included isoform.

**Figure 4.**
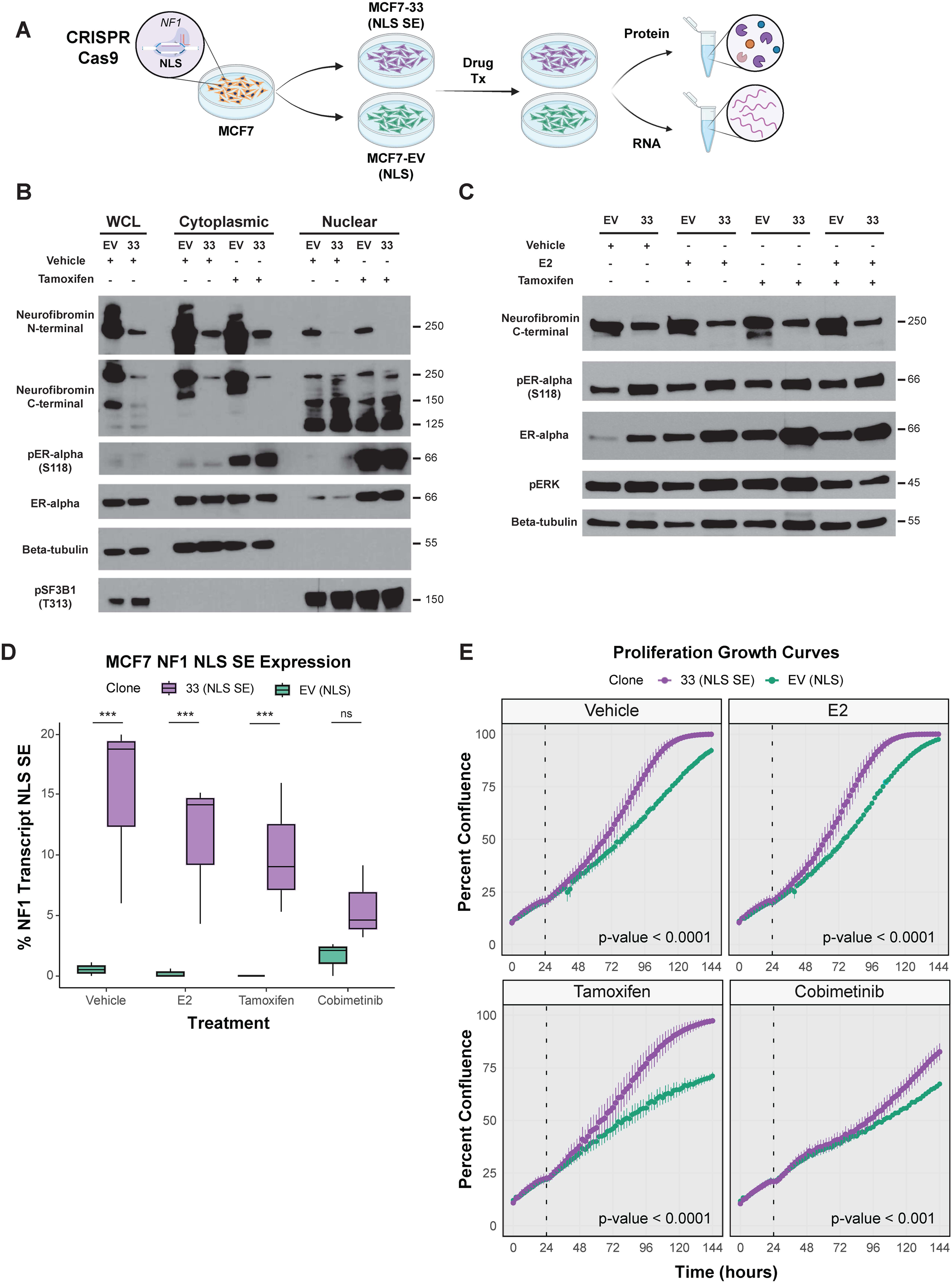
NF1 NLS exon skipping disrupts nuclear localization and enhances proliferative and treatment-resistant growth in ER⁺ breast cancer cells. (A) Schematic of the CRISPR/Cas9 editing strategy used to generate *NF1* NLS exon–skipped mutant MCF7-C33 cells. A single guide RNA (sgRNA) targeting the 5′ end of the NLS exon promoted stable expression of the *NF1* NLS skipping isoform. (B) Western blot analysis of nuclear and cytoplasmic fractions from empty vector (EV) and MCF7-C33 cells treated with vehicle or tamoxifen. Nuclear neurofibromin was absent in MCF7-C33 cells, while cytoplasmic expression was retained, confirming loss of nuclear localization. (C) rMATS analysis from RNA-seq of EV and MCF7-C33 cells following treatment. MCF7-C33 cells exhibited consistently elevated *NF1* NLS skipping across all conditions except under MEK inhibition (\*\**FDR* < 0.001; ns, not significant). (D) Western blot analysis of whole-cell lysates following treatment with vehicle, 17β-estradiol (E2), or tamoxifen revealed increased phosphorylation of ERα at S118 and elevated pERK levels in MCF7-C33 cells compared to EV controls, indicating enhanced ERK-mediated ERα activation. (E) Real-time proliferation of EV and MCF7-C33 cells treated with vehicle, E2, tamoxifen, or cobimetinib. Cells were monitored for 144 hours by IncuCyte live-cell imaging, with treatments added at 24 hours (vertical dashed line). Linear mixed-effects modeling revealed a significant Group × Treatment × Time interaction (*F* = 30.7), indicating distinct growth trajectories. Post hoc analysis confirmed significantly increased proliferation of MCF7-C33 cells under all conditions (vehicle, E2, tamoxifen, *p* < 0.0001; cobimetinib, *p* < 0.001). These findings demonstrate that *NF1* NLS exon skipping promotes estrogen-independent growth and resistance to endocrine and MAPK-targeted therapies.

To confirm loss of nuclear localization, we performed subcellular fractionation followed by Western blotting of nuclear and cytoplasmic lysates. MCF7-C33 cells showed complete loss of nuclear neurofibromin with retained cytoplasmic expression, confirming that NLS SE abolishes nuclear localization (Figure 4B). This phenotype persisted under both vehicle and tamoxifen treatment, indicating that loss of nuclear neurofibromin is a stable feature of the edited cells. Subsequent validation of NF1 NLS SE expression was conducted through RNA sequencing and rMATS analysis of MCF7 clones treated with vehicle, estrogen stimulation (E2), selective estrogen receptor modulator (SERM) (Tamoxifen), or MEK inhibitor (Cobimetinib). MCF7-C33 expressed enriched levels of NF1 NLS SE transcripts compared to EV clones (Figure 4C). Given that our gene set enrichment analysis (GSEA) of the AURORA dataset indicated altered estrogen and RAS signaling, we conducted Western blot analysis of whole-cell lysates from EV and MCF7-C33 cells treated with vehicle, estrogen stimulation, tamoxifen, or a combination of estrogen stimulation and tamoxifen. Western blot results demonstrated that in the absence of neurofibromin nuclear localization, MCF7-C33 cells exhibited increased ER-alpha expression and activation with slight elevated expression in downstream RAS signaling, as indicated by phospho-ER-alpha and phospho-ERK levels (Figure 4D). To determine whether NF1 NLS SE confers a proliferative advantage under therapeutic stress, we performed live-cell imaging using the IncuCyte system. Growth kinetics were modeled using a linear mixed-effects approach incorporating group, treatment, and time as fixed effects. MCF7-C33 cells exhibited significantly higher proliferation rates than EV controls across all conditions, including vehicle (p < 0.0001), E2 (p = 0.0001), tamoxifen (p < 0.0001), and cobimetinib (p < 0.0001) (Figure 4E). These data demonstrate that loss of nuclear neurofibromin enhances ER-alpha and RAS-ERK signaling, promotes estrogen-independent proliferation, and confers resistance to both endocrine and MAPK-targeted therapies. This in vitro model further supports that NF1 NLS SE drives oncogenic adaptation by decoupling neurofibromin nuclear tumor-suppressive function from cytoplasmic RAS regulation.

### Integrative transcriptomic analysis reveals shared oncogenic signaling across NF1 NLS SE models

To further delineate the impact of NF1 NLS SE expression across breast cancer models and clinical datasets, we examined the landscape of differentially expressed genes using an UpSet plot to compare gene level changes across treatment conditions in MCF7 clones as well as metastatic tumors with increased NF1 NLS SE expression from the AURORA cohort (Figure 5A-B). Surprisingly, there was minimal overlap in differentially expressed genes across all datasets, highlighting a high degree of context specificity and potential compensatory signaling dependent on treatment condition or tumor context. Despite the lack of shared gene level expression changes, we hypothesized that convergent biological processes may still be dysregulated. To test this, we performed gene set enrichment analysis across the datasets and identified several common cancer hallmark pathways enriched in NF1 NLS SE upregulated contexts (Figure 5C). Notably, the estrogen response early and late gene sets were consistently downregulated, suggesting impaired ERα driven transcriptional programs in the absence of nuclear neurofibromin. Conversely, pathways associated with inflammatory response, TNFα signaling via NFκB, and KRAS signaling were upregulated across most datasets. These findings underscore a model in which NF1 NLS exon skipping diminishes nuclear neurofibromin function, leading to both loss of ERα transcriptional activity, despite elevated expression of phosphorylated ERα, and activation of pro-tumorigenic signaling cascades. The upregulation of KRAS and inflammatory pathways, which are implicated in cancer cell survival, immune evasion, and therapeutic resistance is especially relevant in ER+ breast cancer, where such pathway activation has been linked to disease progression and metastasis. Additionally, the concurrent downregulation of estrogen responsive genes may reflect a shift towards endocrine resistance, further implicating NF1 NLS SE in driving poor clinical outcomes in Luminal subtypes.

**Figure 5.**
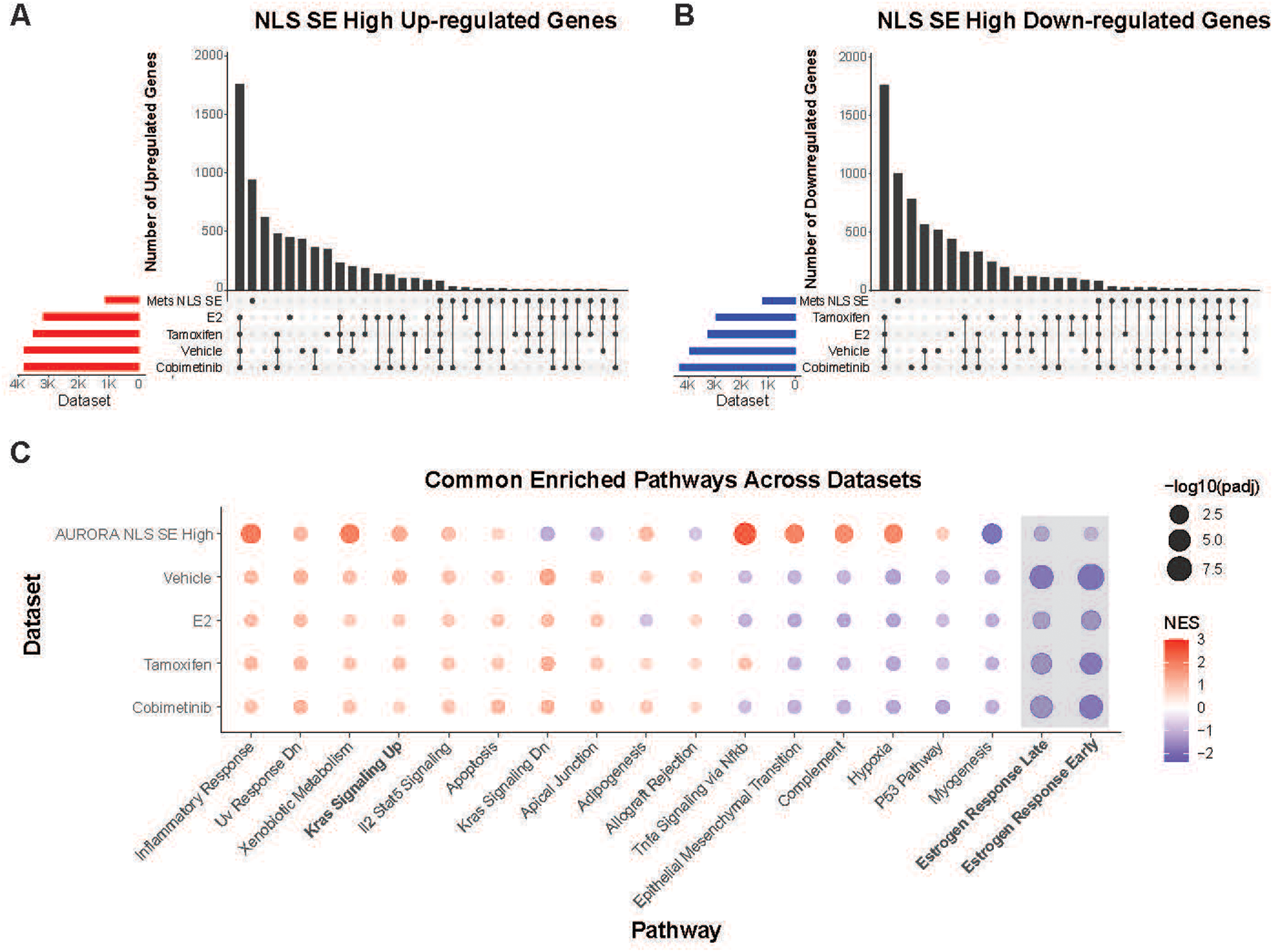
NF1 NLS exon skipping drives convergent transcriptional reprogramming across patient tumors and in vitro models. (A–B) UpSet plots depicting the number of significantly upregulated (A) and downregulated (B) genes across datasets derived from metastatic tumors with increased *NF1* NLS skipping (AURORA cohort; FDR < 0.10; |log₂FC| > 1) and from MCF7-derived clones treated with vehicle, 17β-estradiol (E2), tamoxifen, or cobimetinib (FDR < 0.05; |log₂FC| > 1). Differential gene expression analyses were performed independently for each dataset, and genes were categorized by directionality of change. (C) Heatmap summarizing enriched Hallmark of Cancer gene sets identified by Gene Set Enrichment Analysis (GSEA) across all datasets. Normalized enrichment scores (NES) and –log₁₀-adjusted *p*-values highlight convergence on shared biological processes, including downregulation of estrogen response pathways and upregulation of KRAS and inflammatory signaling networks.

### Loss of Canonical ERα Transcriptional Activity Suggests Non-Genomic Functions Drive Splicing Mediated Resistance in NF1 NLS SE Models

While western blot analysis in Figure 4D demonstrated elevated ERα protein levels in NF1 NLS SE expressing MCF7 clone, gene expression analysis in Figure 5 revealed paradoxical downregulation of canonical estrogen response pathways, including both early and late estrogen response genes. This apparent discrepancy between ERα protein abundance and decreased transcriptional output of estrogen response target genes prompted us to hypothesize that in the absence of nuclear neurofibromin, ERα may be functioning non-canonically as an RBP. The RNA binding capability of ERα is a recently recognized function that we hypothesize is highly relevant to breast cancer progression. Specifically, we speculated that ERα may contribute to splicing regulation to promote survival and resistance mechanisms. To test this, we conducted an RNA pull down assay using oligo(dT) probes to isolate polyadenylated transcripts and assess interactions between ERα and RNA after 24 and 48 hours of treatment (Figure 6A-C). Our results demonstrated increased RNA-bound ERα in NF1 NLS SE expressing MCF7 clone, especially following estrogen and tamoxifen treatment. Notably, we observed an elevation in phosphorylated SF3B1 (pSF3B1 T313), a key spliceosome component and marker of active splicing, suggesting that NF1 NLS SE expression leads to enhanced RNA splicing activity^48^. To further dissect these splicing changes, we performed rMATS analysis to identify differentially spliced events between EV and NF1 NLS SE MC7 clones across treatment conditions (Figure 6D). This analysis revealed a heightened burden of alternative splicing events in NF1 NLS SE expressing clones, consistent with the observed increase in splicing machinery activation. To identify the RBPs potentially driving these splicing changes, we performed motif enrichment analysis using rMAPS2 on the rMATS derived splice junctions. Our analysis revealed a distinct pattern of RBP motif enrichment that varied by treatment and genotype. NF1 NLS SE expressing clones treated with E2 and tamoxifen exhibited enriched splicing activity associated with BRUNO4/6 (also known as CELF proteins). These RBPs are known to regulate NF1 exon 31 splicing within the GRD, directly linking them to RAS signaling regulation. Enrichment of ESRP1 motifs, a key regulator of epithelial-to-mesenchymal transition (EMT) splicing programs and poor prognosis marker in ER+ breast tumors, was also observed, aligning with our prior EMT gene expression findings^49–52^. Additional RBPs motifs enriched under these treatments included: RBM8A, which supports breast cancer progression via IGF1R-PI3K signaling, FMR1, linked to oncogenic transformation in multiple cancers, and FUS, implicated in breast cancer liver metastasis through splicing of circular EZH2 RNA^53^. Cobimetinib treated NF1 NLS SE expressing cells exhibited strong enrichment for SRSF1 and SRSF9 motifs, both of which are well-established proto-oncogenic splicing regulators that drive tumorigenesis and metastatic behavior^11^. Collectively, these data suggest that loss of nuclear neurofibromin enables ERα and other RBPs to reprogram splicing networks to promote transcriptional plasticity, survival, and resistance in ER-positive breast cancer. This shift toward RBP-driven transcriptome remodeling highlights a novel mechanism of endocrine therapy evasion linked to NF1 alternative splicing.

**Figure 6.**
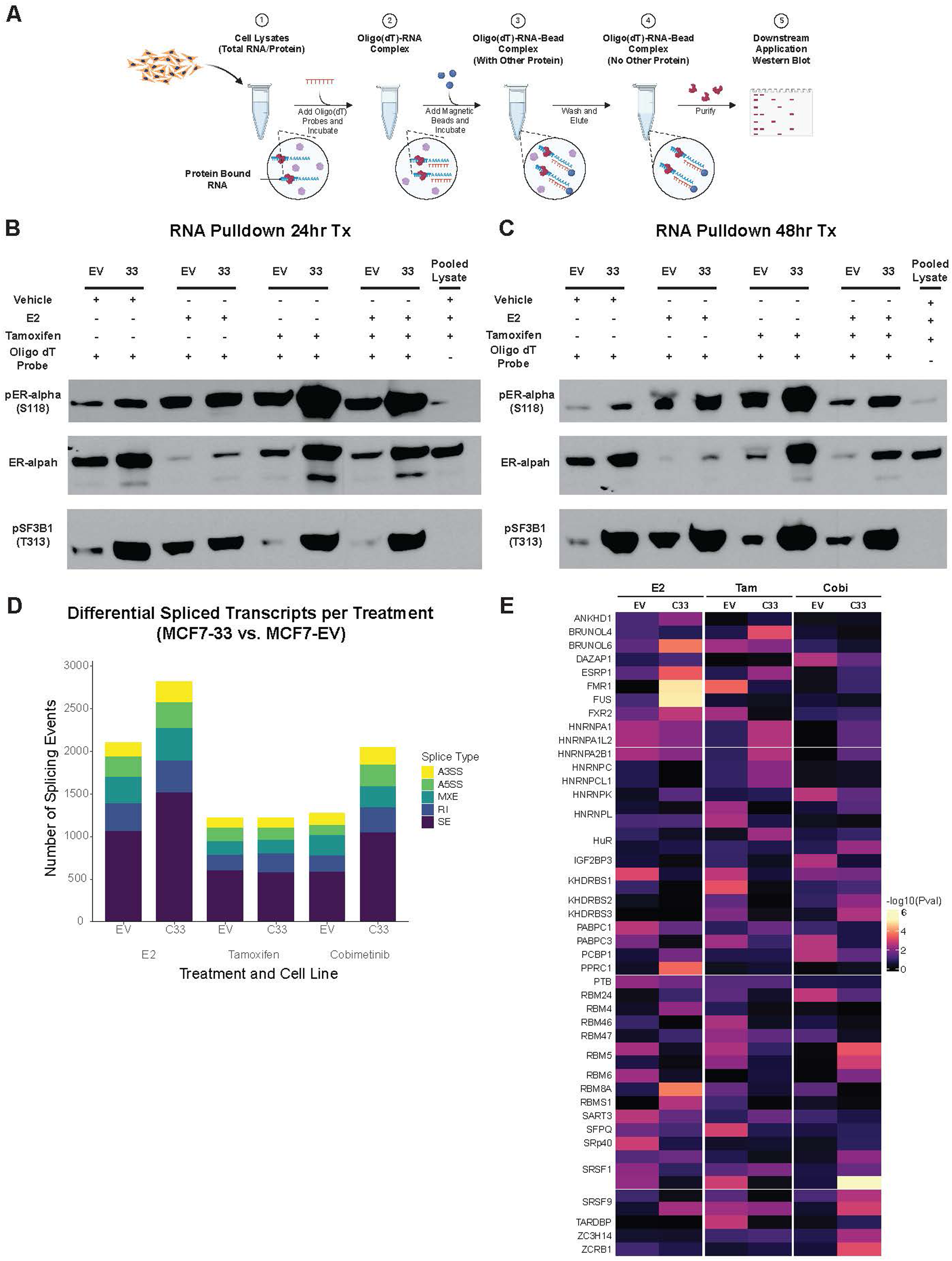
NF1 NLS exon skipping enhances RNA-bound ERα and splicing activity in luminal breast cancer cells. (A) Schematic of the oligo(dT)-based RNA pulldown workflow. Polyadenylated mRNAs were isolated from total cell lysates using oligo(dT) probes, and RNA–protein complexes were captured with magnetic beads for downstream Western blot analysis. (B–C) Western blot analysis of RNA-bound protein fractions from MCF7 empty vector (EV) and MCF7-C33 cells following 24-hour (B) or 48-hour (C) treatment with vehicle, 17β-estradiol (E2), or tamoxifen. MCF7-C33 cells exhibited increased RNA-associated ERα and enhanced splicing factor binding compared to EV controls. (D) Quantification of differentially spliced events identified by rMATS between EV and MCF7-C33 cells under vehicle, E2, tamoxifen, or cobimetinib treatment. Events were categorized by splicing type: skipped exon (SE), retained intron (RI), mutually exclusive exon (MXE), alternative 3′ splice site (A3SS), and alternative 5′ splice site (A5SS). (E) Heatmap of RNA-binding protein (RBP) motif enrichment from rMAPS2 analysis of splice junctions identified in panel D. Enriched motifs across treatment conditions highlight differential RBP activity and altered splicing regulation in *NF1* NLS SE–expressing cells.

## Discussion

In this study, we identify a previously unrecognized role for *NF1* alternative splicing in breast cancer progression and establish a functional link between the *NF1* NLS-skipped isoform (NLS SE) and endocrine-resistant, ER-positive disease. We show that NLS SE is enriched in metastatic tumors, occurs independently of *NF1* genomic alterations, and is associated with poor overall survival, particularly in luminal breast cancer^16^. These findings position *NF1* NLS SE as both a biomarker of aggressive disease and a mechanistic driver of therapeutic resistance^54^.

Importantly, our results demonstrate that *NF1* transcript dysregulation represents a distinct regulatory axis separate from canonical genomic loss^3,55^. While prior studies have focused on *NF1* mutations and their effects on RAS signaling, our data reveal that splicing-mediated loss of the NLS disrupts nuclear neurofibromin localization and function^29^. This loss alters the balance of ERα and RAS signaling, decoupling nuclear tumor-suppressive activity from cytoplasmic signaling control^34^. Notably, despite increased ERα protein abundance and activation, canonical estrogen-responsive transcriptional programs were suppressed, suggesting a fundamental shift in ERα function^14,56^.

We propose that, in the absence of nuclear neurofibromin, ERα adopts a non-canonical role as an RNA-associated factor that contributes to splicing regulation. Consistent with this model, we observed increased RNA-bound ERα and enhanced phosphorylation of SF3B1, indicative of heightened spliceosome activity. Transcriptomic and motif enrichment analyses further identified activation of RNA-binding protein networks implicated in tumor progression and endocrine resistance, including CELF family proteins, ESRP1, and SRSF family members^57^. These findings suggest that *NF1* NLS SE promotes a reprogrammed, RBP-driven transcriptomic landscape that supports tumor cell survival under therapeutic pressure.

Our data also highlight the plasticity of this regulatory network across treatment contexts. Distinct RBP motifs were enriched following estrogen stimulation, tamoxifen treatment, or MEK inhibition, indicating that splicing reprogramming is dynamically shaped by signaling environment. This adaptability may underlie the ability of tumor cells to maintain growth and survival despite targeted therapeutic interventions.

From a translational perspective, *NF1* NLS SE represents a clinically relevant isoform-level biomarker that may improve risk stratification in ER-positive breast cancer. Given its association with poor prognosis and resistance to both endocrine and MAPK-targeted therapies, targeting splicing regulatory networks or restoring nuclear neurofibromin function may represent promising therapeutic strategies. These findings further underscore the need to incorporate transcript-level analyses into the characterization of tumor suppressor dysregulation.

While our findings demonstrate a strong association between NF1 NLS SE, increased RNA-bound ERα, and widespread splicing alterations, several limitations should be considered. First, although ERα-RNA association was consistently elevated across experimental conditions, we did not directly assess transcript-specific binding or perturb ERα RNA-binding activity to establish causality in splicing regulation. Second, the magnitude of exon skipping observed in patient samples was modest, though such changes may have outsized functional consequences given the role of the NLS in regulating neurofibromin localization. Finally, while our in vitro model recapitulates key signaling and proliferative phenotypes, in vivo validation will be necessary to fully define the contribution of NF1 splicing to tumor progression and therapeutic resistance. Future studies addressing these questions will further refine the mechanistic framework proposed here.

Finally, recent structural studies demonstrating that neurofibromin functions as a homodimer raise important questions regarding the impact of isoform diversity on protein assembly and function. Loss of the NLS-containing isoform may alter dimer composition, stability, or subcellular distribution, potentially generating functionally distinct complexes. Future studies dissecting how *NF1* isoform balance influences dimerization, interaction networks, and nuclear signaling will be critical for understanding how splicing-dependent structural changes contribute to tumor progression^30,58,59^. Elucidating these mechanisms may uncover new therapeutic vulnerabilities in cancers characterized by *NF1* splicing dysregulation.

## Methods

### Access and Processing of AURORA and TCGA RNA-seq Data

Access to RNA sequencing data from the AURORA US Metastatic Breast Cancer Project (phs002611.v1.p1) and The Cancer Genome Atlas (TCGA; phs000178.v11.p8) was obtained through authorized access via the NCBI database of Genotypes and Phenotypes (dbGaP). Data access requests were approved by the relevant Data Access Committees and managed in accordance with NIH Genomic Sharing (GDS) policy. Following approval, raw sequencing files, FASTQ, were retrieved using the SRA Toolkit (v3.0.0), which allows for secure download and decryption of controlled-access sequencing data. All data were stored and processed within an institutional secure computing environment compliant with dbGaP security Guidelines. All file handling, storage, and downstream analysis followed protocols ensuring confidentiality, access control, and audit compliance as stipulated by the NIH Data Use Certification Agreement.

### Alternative Splicing Analysis Using rMATS

*NF1* isoform usage between primary and metastatic breast tumors were identified using the differential splicing analysis rMATS (v4.1.0), a robust statistical tool designed to detect changes in alternative splicing from RNA-seq data. Aligned BAM files from each group (primary vs. metastatic) were organized into two input lists and analyzed using paired-end mode. The reference genome annotation was provided using a GTF file based on GENCODE v38, and splicing was assessed across known and novel junctions. The analysis was configured to account for variable read lengths. For rMATS analysis, splicing events with FDR < 0.05 and a minimum of 5 junction spanning reads were considered significant. Events were further prioritized if they were observed across multiple patients or involved functionally relevant regions of the *NF1* transcript.

### Survival Analysis

Overall survival analyses were performed using clinical and genomic data from the TCGA Breast Cancer cohort. Samples with missing data for survival status, survival time, age at diagnosis, mutation burden, PAM50 subtype, or *NF1* NLS SE status were excluded. Splicing classification was based on exon-level quantification from TCGA SpliceSeq, with tumors categorized as either *NF1* NLS SE or *NF1* NLS Inclusion. Survival time was right-censored at 120 months to define a 10-year survival endpoint. Kaplan-Meier estimates were generated using the survfit function and visualized with ggsurvfit. Univariable and multivariable Cox proportional hazards models were fit using coxph function. The multivariable model included *NF1* NLS spicing status, age at diagnosis, mutation count, *NF1* status, and *TP53/PIK3CA* mutation status. To account for intrinsic subtype, a stratified Cox model was used with PAM50 as a stratification factor. Proportional hazards assumptions were tested with Schoenfeld residuals. Results were summarized using gtsummary, reporting hazard rations, 95% confidence intervals, and p-values, with significance set at p < 0.05.

### Generation of *NF1* NLS SE Human Breast Cancer Model

To evaluate the effect of *NF1* NLS SE expression in ER+ human breast cancer, we utilized the ER+ MCF7 human adenocarcinoma cell line. Using CRISPR/Cas9 technologies, we targeted exon 52 of 58 that encodes the NLS within the C-terminal domain of *NF1*. We established a cell line that expresses stable levels of *NF1* NLS SE, cell line MCF7-C33, and chose it for further analysis. The MCF7-C33 cell line contains a heterozygous insertion of a thymine in *NF1* that introduces a premature stop codon. Exon skipping of the NLS removes the premature stop codon allowing for translation of *NF1* NLS SE isoforms.

### Cell Culture

Human ER+ MCF7 breast cancer cells (HTB-22) were cultured in phenol red-free Dulbecco’s Modified Eagle Medium (DMEM/F12; Gibco, Cat# 21041025) supplemented with 10% fetal bovine serum (FBS; VWR, Cat# 1300-500) and 5 mL penicillin-streptomycin (Gibco; Cat# 15140122). All cell lines were maintained at 37°C in a humidified incubator with 5% CO_2_. Routine mycoplasma testing was performed every six months using the ATCC Universal Mycoplasma Detection Kit (Cat# 30-1012-K) and confirmed all cultures were free of contamination.

### IncuCyte Proliferation Analysis

Growth curves for MCF7-EV and MCF7-C33 clones were analyzed under treatment of ethanol (Vehicle), 10 nM β-estradiol, 100 nM tamoxifen (Sigma), or 100 nM cobimetinib (Genentech) for 144 hours using Sartiorius IncuCyte imaging. Treatments wre added 24 hours post-plating, and cell confluency was recorded every 2 hours. A linear mixed-effects model was fit with confluency as the response variable. The model included fixed effects for Group (EV vs. C33), Treatment, and Time (continuous, hours), and all interaction terms, and a random intercept for Time to account for repeated measurements. Type III ANOVA was conducted with Kenward-Roger approximation to test the significance of fixed effects. Post hoc pairwise comparisons between clones within each treatment were performed, with Tukey-adjusted p-values to control for multiple comparisons. Model residuals were checked for normality and homoscedasticity. Interaction terms (Group x Treatment x Time) were included a priori based on hypothesized differential treatment responses and were interpreted as growth rate differences between clones. Significance was set a p < 0.05.

### RNA Sequencing

RNA sequencing was conducted on MCF7 Clones using three biological replicates per condition. For each replicate, 1.2 x 10^5^ cells were seeded into individual wells of a 6-well plate (Corning, Cat# 3506) and treated with ethanol (Vehicle), 10 nM β-estradiol, 100 nM tamoxifen (Sigma), or 100 nM cobimetinib (Genentech) for 24 hours. After 24 hours of treatment, cells were lysed directly with RLT buffer (RNeasy Kit, Qiagen, Cat# 74104). Lysates were transferred to QIAshredder columns (Qiagen, Cat# 79654). An equal volume of 70% ethanol (Pharmco, Cat# 111000200) was added to the flow-through and applied to RNeasy spin columns for RNA purification following Manufacturer’s protocol. RNA library preparation and sequencing were performed by the Van Andel Genomics Core. Libraries were prepared using 500 ng of total RNA with KAPA mRNA HyperPrep Kit (Kapa Biosystems, v4.17). RNA was enzymatically fragmented to 300-400 bp, and cDNA libraries were ligated to IDT for Illumina TruSeq Unique Dual (UD) Indexed adapters. Library quality was assessed using Agilent DNA High Sensitivity chip, the QuantiFluor® dsDNA system (Promega), and KAPA qPCR-based library quantification. Indexed libraries wre pooled and sequenced on an Illumina NovaSeq 6000 platform using 200-cycle S4 kit, generating 100 bp paired-end reads at an approximate depth of 70 million reads per sample. Base calling was performed using Illumina RTA3, and sequencing output was demultiplexed and converted to FASTQ format using Illumina Bcl2fastq v1.9.0. Adapter trimming was conducted with Trim Galore (v0.6.0). Reads were aligned to GRCh38 reference genome (GENCODE release v38) using STAR (v2.7.8a) in two-pass basic mode.

### Cell Fractionation

To isolate nuclei for downstream applications, cells were subjected to a two-step lysis and centrifugation protocol. Cells were first resuspended in ice-cold Cell Lysis Buffer (20 mM Tris-HCL, 85 mM KCl, 0.5% NP-40) and incubated on ice for 15 minutes, with gentle vortexing every 5 minutes. Following lysis, the suspension was transferred to a pre-chilled dounce homogenizer and mechanically disrupted using 10 strokes with a tight-fitting pestle. The homogenate was transferred to a 1.5 mL microcentrifuge tube and centrifuged at 800 x g for 5 minutes at 4°C. The supernatant was carefully aspirated, and the nuclear pellet was rinsed by gently dispensing 1 mL of ice-cold Cell Lysis Buffer. The nuclei were pelleted by centrifugation at 1,700 x g for 5 minutes at 4°C. The wash step was repeated once more to ensure removal of cytoplasmic contaminants. Finally, the nuclear pellet was resuspended in Shearing Buffer (10mM Tris-HCl pH 7.6, 1 mM EDTA, 0.1% SDS) and processed immediately for Western blot analysis or stored at −80°C for future use.

### RNA Pulldown

Polyadenylated mRNAs were isolated from MCF7 lysates using the Sigma-Aldrich Dynabeads® mRNA DIRECT™ Purification Kit (Cat. No. 61011), following manufacturer’s protocol. Specifically, steps 2 through 6 of the kit protocol were performed to prepare and sell Oligo(dT) magnetic beads, allowing for the capture of mRNA via hybridization to poly(A) tails. For lysate preparation, MCF7 cells were washed twice with ice-cold PBS and scraped into ice-cold lysis buffer [20mM Tris-HCl (pH 7.4), 250 mM NaCl, 10 mM KCl, 5 mM MgCl_2_, 0.1% Triton X-100, RNasin® Plus RNase Inhibitor (Promega), 1x protease inhibitor and phospho-protease inhibitor cocktail]. Samples were incubated on ice for 10 minutes and vortexed every 5 minutes. Cell debris was removed by centrifugation at 14,000 x g for 10 minutes at 4°C, and the clarified supernatant was collected. Protein concentration was measured using a Bradford Protein Assay (Bio-Rad), and equal amounts of protein lysates were incubated with pre-swollen Oligo(dT) magnetic beads for 2 hours at 4°C with end-over-end rotation to allow mRNA-protein complex capture. After incubation, samples were centrifuged at 2,500 x g for 2 minutes to pellet the magnetic beads. Beads were washed five times with the provided Wash Buffer, and mRNA-protein complexes were eluted in 2x Laemmli buffer supplemented with 10 mM DTT. Eluates were boiled at 95°C for 10 minutes, with vortexing every minute. Proteins associated with polyadenylated mRNA were then analyzed by SDS-PAGE followed by Western blotting.

### Western Blots

Cells were cultured in their designated growth media and treated with ethanol (Vehicle), 10 nM β-estradiol, 100 nM tamoxifen (Sigma), or 100 nM cobimetinib (Genentech) for 24 hours prior to lysate preparation. Following treatment, cells were rinsed with ice-cold 1X PBS (Gibco, Cat# 10010023), scraped, and lysed using Rb lysis buffer (20 mM Tris-HCl, pH 7.6; 5 mM EDTA; 150 mM NaCl; 0.5% NP-40; 50 mM NaF; 1 mM β-glycerophosphate), supplemented with PhosStop phosphatase inhibitors (Roche, Cat# 04906837001) and a protease inhibitor cocktail (Sigma, Cat# 11873580001). Protein samples were subsequently separated by SDS-PAGE. Immunoblotting was carried out using antibodies against neurofibromin – Bethyl (Bethyl, Cat# A300-140A, primary 1:500, secondary 1:2000), H12 (Santa Cruz Biotechnology, Cat# sc-376886, primary 1:500, secondary 1:2000), pERα-S118 (abcam Cat# ab32396, primary 1:1000, secondary 1:2000), ERα (Cell Signaling Cat# 8644, primary 1:1000, secondary 1:2000), Phospho-SF3B1 T313 (Cell Signaling Cat#25009, primary 1:1000, secondary 1:2000), β-Tubulin (Cell Signaling Cat#2146, primary 1:2000, secondary 1:2000), phosphor-ERK (Cell Signaling Cat#4370, primary 1:1000, secondary 1:2000).

### Statistical Analysis

All statistical analyses were performed in R (version 4.2.2). Linear mixed-effects were fit using the ‘lme4’ and ‘emmeans’ packages to assess proliferation over time across cell lines and treatments. These models included fixed effects for Group, Treatment, Time, and random intercepts for Time. ANOVA tests were performed using type III sums of squares, and post hoc pairwise contrasts were adjusted using Tukey’s method. Survival analysis was conducted using Kaplan-Meier estimates and Cox proportional hazard regression. Multivariable models included covariates for mutation count, diagnosis age, *NF1* status, *TP53* and *PIK3CA* mutation status, and were stratified by PAM50 subtype. Model assumptions were verified using Schoenfeld residuals. *NF1* splicing-to-mutation association was tested using Fisher’s exact tests and multivariable logistic regression. Splicing differences between matched primary and metastatic tumors were evaluated using a beta-binomial generalized linear model (GLM) to account for matched samples from patients with multiple metastatic lesions. To assess associations between *NF1* NLS SE expression and genomic alterations, splicing data from the TCGA Breast Cancer Cohort were obtained from the TCGA SpliceSeq database. Genomic status for genes were retrieved from cBioPortal.

## Code and Data Availability

Computational analyses performed in this study are available through a public GitHub repository: https://github.com/dischip/NF1_NLS_BreastCancer_Manuscript. This repository contains R scripts, metadata files, session information, and environmental configuration files required to reproduce survival analyses, differential expression modeling, splicing association testing, and proliferation assay statistics. Raw sequencing data were not uploaded due to side and access limitations but are available from TCGA and the AURORA consortium under controlled access.

## Author Contributions

PD conceived and designed the study. PD and SA performed experiments. PD, SA, and MA acquired data. PD, SA, IB, and EW analyzed and interpreted the data. PD drafted the manuscript, and CG and MS provided critical revisions. All authors reviewed and approved the final version of the manuscript.

## Acknowledgments

We are grateful to Dr. Hui Shen, Dr. Tim Triche, Dr. Yi Xing, and Dr. Olga Anczuków for their thoughtful discussions, guidance, and critical feedback that helped shape this work. We also thank the Van Andel Institute Genomics Core (RRID:SCR_022913) for sequencing support and the Biostatistics and Bioinformatics Core (RRID:SCR_024762) for their expertise and assistance with data processing, analysis, and interpretation.

